# Identification of a conserved virion-stabilizing network inside the interprotomer pocket of enteroviruses

**DOI:** 10.1101/2020.07.06.189373

**Authors:** Justin W. Flatt, Aušra Domanska, Alma L. Seppälä, Sarah J. Butcher

## Abstract

Major efforts have been underway to develop broad-spectrum high potency capsid binders that inhibit the life cycle of enteroviruses, a large group (family *Picornaviridae*) whose members include poliovirus, coxsackieviruses, echoviruses, numbered enteroviruses, and rhinoviruses. These diverse viruses cause a wide variety of illnesses, ranging from the mild common cold to hand-foot-and-mouth disease, myocarditis, pancreatitis, aseptic meningitis, and encephalitis. So-called classical capsid binders target a surface exposed hydrophobic pocket in one of the viral coat proteins (VP1) to prevent the genome uncoating process. However, efficacy, toxicity, emergence of drug-resistant viruses, and existence of certain enteroviral species that lack the VP1 pocket limit their clinical benefit. Recently, we identified a new druggable site at a conserved interface formed by multiple capsid proteins, the VP1-VP3 interprotomer pocket. To further study the properties that confer druggability at this site, we have determined high-resolution cryo-electron microscopy structures of two enteroviruses, coxsackieviruses B3 and B4, complexed with interprotomer-targeting compounds, CP17 and CP48 respectively. Until now, there has been no structure available for Coxsackievirus B4 despite the fact that the virus has long been implicated in the development of insulin-dependent diabetes mellitus. At better than 3 Å resolution, we could identify the detailed interactions that facilitate ligand binding. Both compounds target the same three conserved residues, each from a different polypeptide chain, to form a virion-stabilizing network inside the pocket. We measured the in silico binding energy for both inhibitors when anchored to the network and found a global stabilizing effect on the order of thousands of kcal/mol under saturating conditions (60 total sites per virion). Intriguingly, a recent X-ray structure has revealed that glutathione targets the same network within the interprotomer site of bovine enterovirus F3, where it is thought to facilitate virus assembly. In summary, our findings provide the structural basis for how a newly designed class of capsid binders target and stabilize enteroviruses. Future efforts to chemically optimize drugs for enhanced targeting to the interprotomer pocket is a promising endeavor in the fight against enteroviruses, especially given the possibility of synergistic effects when used in combination with classical VP1 binders like pleconaril.

## Introduction

The group B coxsackieviruses (CVBs) are a major source of both acute and chronic diseases in humans. Age and immune status are thought to be the main determinants of morbidity and mortality, with infants, young children, and immunocompromised individuals being particularly susceptible to serious and sometimes life-threatening infections. Coxsackievirus B3 (CVB3) can cause cardiac arrhythmias and acute heart failure (Kearney et al., 2001, Massilamany et al., 2014). Additionally, CVB3 infections during pregnancy have been linked to an increase in neurodevelopmental delays, fetal myocarditis, and spontaneous abortions (Euscher et al., 2001, Ornoy and Tenenbaum, 2006). Coxsackievirus B4 (CVB4) appears to elicit or enhance certain autoimmune disorders such as type 1 diabetes as the virus has been isolated from individuals diagnosed with rapid onset type 1 diabetes, and these isolates were then shown to cause diabetes in mice models (Notkins et al., 1979, See and Tilles, 1995). Dotta et al. have provided arguably the most direct support for CVB4 as a viral trigger of diabetes via immunohistochemical detection and sequencing of virus from the pancreatic tissue of diabetic patients (Dotta et al., 2007). Thus, it is of great importance to develop antiviral drugs and vaccines to combat CVBs, as well as other enteroviruses, given that cases and outbreaks can result in substantial hospitalization and burden of healthcare services.

CVB capsids share a common enteroviral architecture constructed from 60 repeating protomers, each consisting of the four structural proteins VP1, VP2, VP3, and VP4 (Jiang et al., 2014). The protomers assemble to form the ~30 nm wide icosahedral shell with a pseudo T = 3 arrangement that encapsidates the linear single-stranded RNA genome. The arrangement occurs because of the similar structures of VP1, VP2, and VP3, which all adopt an eight-stranded, antiparallel β-barrel fold despite having low sequence homology. The four strands of the β-sheets are connected by hypervariable loops that are responsible for the high antigenic diversity of enteroviruses. The organization of the 180 β-barrels is much the same as observed in T = 3 lattices formed by 180 identical copies of a capsid protein, with VP1 localized to 5-folds, while VP2 and VP3 alternate around the 2- and 3-fold axes. VP4 is located on the inside of the capsid and is myristoylated. Many picornaviruses utilize a canyon-like feature on their surface to bind cellular receptors belonging to the immunoglobulin superfamily (Lin et al., 2009). Binding into the canyon destabilizes virions and initiates the uncoating process by triggering release of the lipid moiety “pocket factor” from the small hydrophobic pocket in VP1 (Wang et al., 2012). Notable exceptions include rhinovirus C and parechoviruses, which do not accommodate a fatty-acid pocket factor (Liu et al., 2016, Domanska et al., 2019, van de Ven et al., 2011).

Small molecules that bind tightly and specifically to conserved capsid features to interfere with virus entry or uncoating are among the most promising strategies for blocking enterovirus infections (Baggen et al., 2018). These molecules, the WIN antiviral compounds, target the VP1 hydrophobic pocket, which has an entrance located at the base of the canyon-like depression surrounding each capsid 5-fold axis (Chen et al., 2008). The site is normally occupied by the pocket factor, however, binding of chemically optimized compounds dislodges the lipid due to the drugs having a much higher binding affinity (Smith et al., 1986). Replacement of the pocket factor with capsid binders provides entropic stabilization by raising the uncoating free energy barrier against thermal or receptor-induced conformational changes (Tuthill et al., 2010, Tsang et al., 2000). In this way, the compounds are able to prevent formation of expanded 135S intermediates or A-particles, which is a required step for genome release. In vitro testing has shown this to be the case for several VP1 pocket binders; they possess high potency and broad-spectrum activity against enteroviruses. However, clinical development has been thwarted because of issues related to efficacy and toxicity, as well as emergence of drug-resistant viruses (Thibaut et al., 2012). Recently, we discovered a second druggable pocket at a conserved VP1-VP3 interprotomer interface in the viral capsid (Abdelnabi et al., 2019). This interface is in a region of the capsid that undergoes quaternary conformational changes to promote disassembly and release of the virion’s genome into the host cell. Synthetic compounds that occupy the interprotomer pocket are inhibitors of a large number of enteroviruses, and act synergistically with inhibitors that target the VP1 pocket.

Here, in an effort to better understand the druggable features of the interprotomer pocket, we have analyzed high-resolution structures of two medically important enteroviruses, coxsackieviruses B3 and B4, complexed with interprotomer-targeting compounds CP17 and CP48 respectively. The structures were determined by cryo-electron microscopy (cryo-EM) to beyond 3 Å resolution, which allowed us to identify the detailed interactions that facilitate drug binding at the VP1-VP3 interface. In addition to modeling the key residues, we also calculated interaction energies for both compounds using in silico methods. We found that both compounds target the same interprotomer side chains, and the energy of interaction is comparable to what has been observed for robust, high-affinity binders of the VP1 hydrophobic pocket. These results taken together help to explain how this new class of drugs interferes with virus uncoating, and indicate that it is worthwhile to focus on developing therapies that include a synergistic combination of binders to potentially improve efficacy, alleviate side effects, and shorten treatment of enteroviral infections.

## Results

### CP17 bound to the interprotomer pocket of CVB3

CP17 is a benzenesulfanomide derivative that potently inhibits the CVB3 Nancy strain in cells (EC_50_ 0.7 ± 0.1 μM) via a direct interaction with the capsid that increases virion thermostability by 1.5 and 2.1 log_10_ TCID_50_/mL at 46°C and 49°C, respectively (Abdelnabi et al., 2019). A 4.0 Å cryo-EM structure of CP17 in complex with CVB3 Nancy (EMD-0103) revealed that the site of binding is located at a conserved VP1-VP3 interprotomer interface, but the low resolution of the map prevented identification of the detailed interactions within the pocket. We reprocessed the raw data (EMPIAR-10199) using RELION 3.0 and the resolution improved to 2.8 Å (Zivanov et al., 2018). The cryo-EM map shows pronounced backbone features for the four structural proteins of CVB3, and well defined density for CP17 on the surface of the capsid (Figure 1A and 1C). Importantly, the resolution is now sufficient for describing specific ligand-protein interactions (Figure 1B). The interprotomer site is located between adjacent asymmetric units “protomers” in a narrow opening formed at the intersection of neighboring VP1 β-barrels and the C-terminus of a proximally situated VP3 molecule. Three residues play a key role in binding CP17 and are conserved across enteroviruses. In CVB3 Nancy they are Arg219 (VP1), Arg234 (VP1), and Gln233 (VP3). The Arg residues, which come from neighboring VP1 polypeptide chains, are situated in the deepest part of the pocket, and in the high-resolution structure, we observed that their guanidinium groups form salt bridges with the carboxylic end of CP17. The Gln residue from the C-terminus of VP3 is positioned at the entrance of the drug site, where the oxygen in the side chain engages in a hydrogen bond interaction with the NH located in the elbow region of the inhibitor. The binding energy (sum of van der Waals and electrostatics) for CP17 is −74 kcal/mol based on the NAMD energy plugin in VMD. The predicted value is comparable to potent inhibitors of the VP1 hydrophobic pocket, namely the capsid binders GPP3 (−66 kcal/mol) and NLD (either −69 or −64 kcal/mol depending on protonation state) (De Colibus et al., 2015).

**Figure 1:**
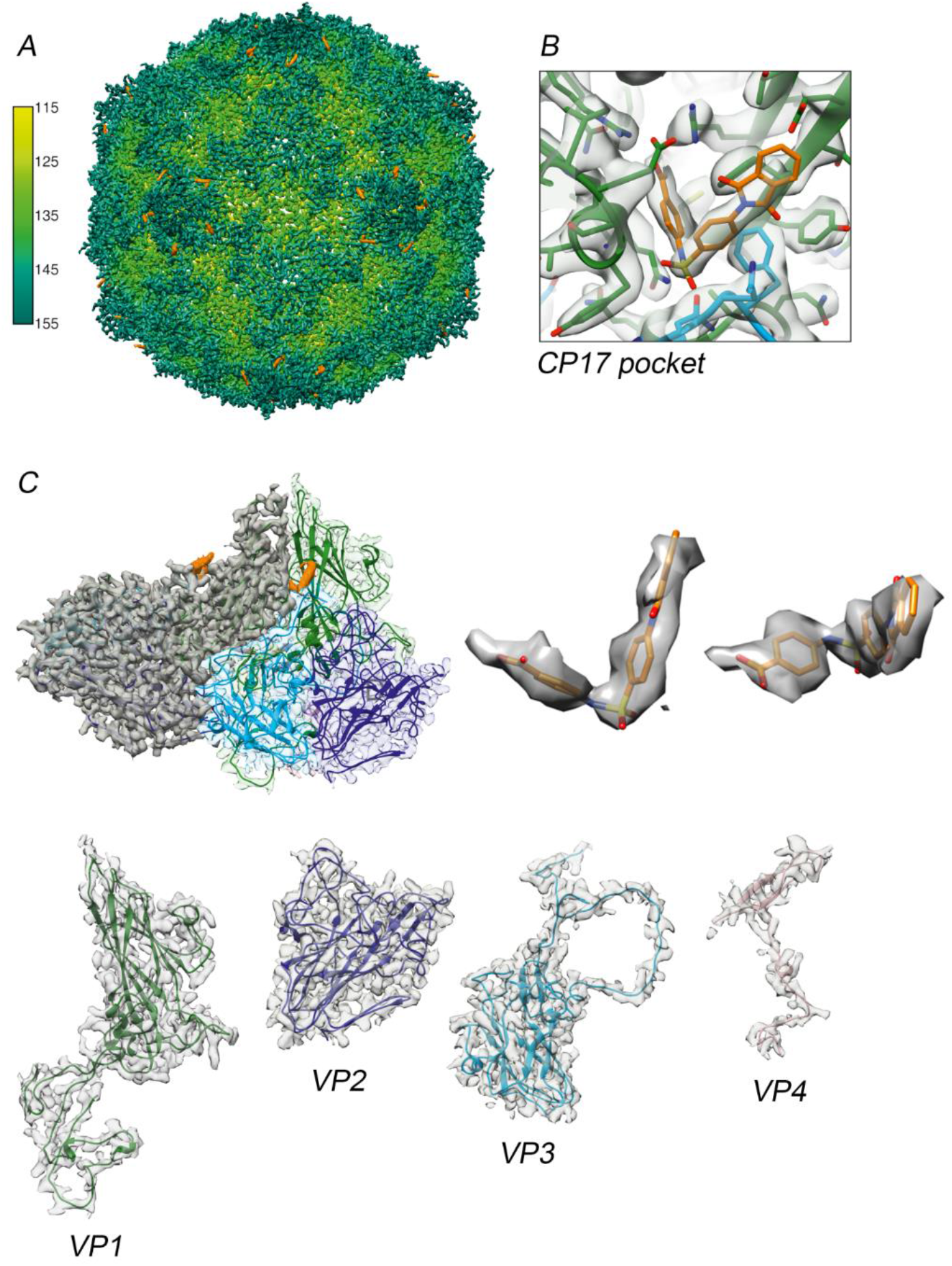
Cryo-EM structure of CP17-bound CVB3. (A) Three-dimensional reconstruction of CVB3 after incubation with a saturating amount of capsid binder. The virion is viewed along the icosahedral 2-fold axis and colored according to radial distance in Å from the particle center. Density for CP17 is shown in orange. (B) A close-up view of the capsid binder within the interprotomer pocket. Both inhibitor and pocket side chains are well resolved. (C) The 2.8 Å resolution map allows unambiguous construction of an atomic model for the capsid proteins and CP17.

### CP48 within the CVB4 virion

To further investigate and confirm the activity and structural basis for how inhibitors bind at the interprotomer pocket, we determined a structure of CVB4 in the presence of a commercially available analog, which we refer to as CP48. This particular inhibitor was found to be active against all six serotypes of CVBs, and completely inhibited poliovirus type 1 replication at a concentration of 144 μM (Abdelnabi et al., 2019). We chose to work with CVB4 because no structure exists for this important human pathogen. First, we confirmed that addition of CP48 increases CVB4 thermal stability (Figure 2).

**Figure 2:**
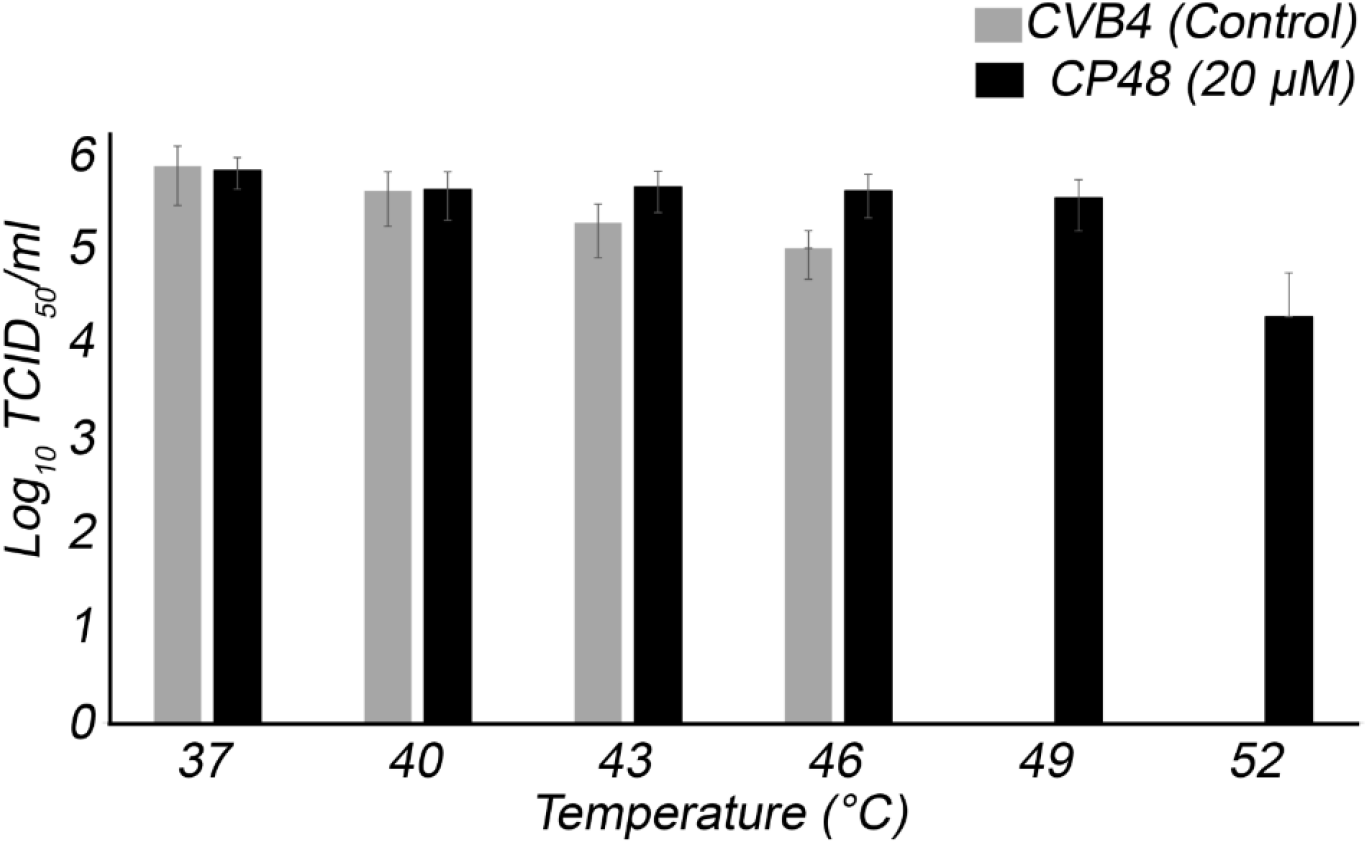
CP48 effects on CVB4 thermal stability. Thermostability assay in the presence (black) or absence (gray) of interprotomer-targeting CP48.

Then, purified virus was incubated with a saturating concentration of CP48 (virus:drug molar ratio of 1:2500), applied to grids, and flash frozen for cryo-EM. After image processing, a subset of 18,626 particles yielded a 2.7 Å reconstruction using the FSC 0.143 threshold criterion (Rosenthal and Henderson, 2003). The outer surface of the virus particle is similar to that of other enteroviral capsids, with major features including the 5-fold star-shaped mesas, 3-fold propeller-like protrusions, and 2-fold depressions (Figure 3A). In addition, there is stable and well defined density for the inhibitor at the interprotomer site (Figure 3B). Modeling confirmed the critical role of the three pocket side chains: two Arg residues on the inner surface and a Gln residue at the entrance. The energy for the bound inhibitor is approximately −45 kcal/mol versus −74 kcal/mol for CP17-CVB3, which the EC_50_ for CP48-CVB4 is only 8.6 ± 0.8 μM compared to an EC_50_ of 0.7 ± 0.1 μM for CP17-CVB3 (Abdelnabi et al., 2019).

**Figure 3:**
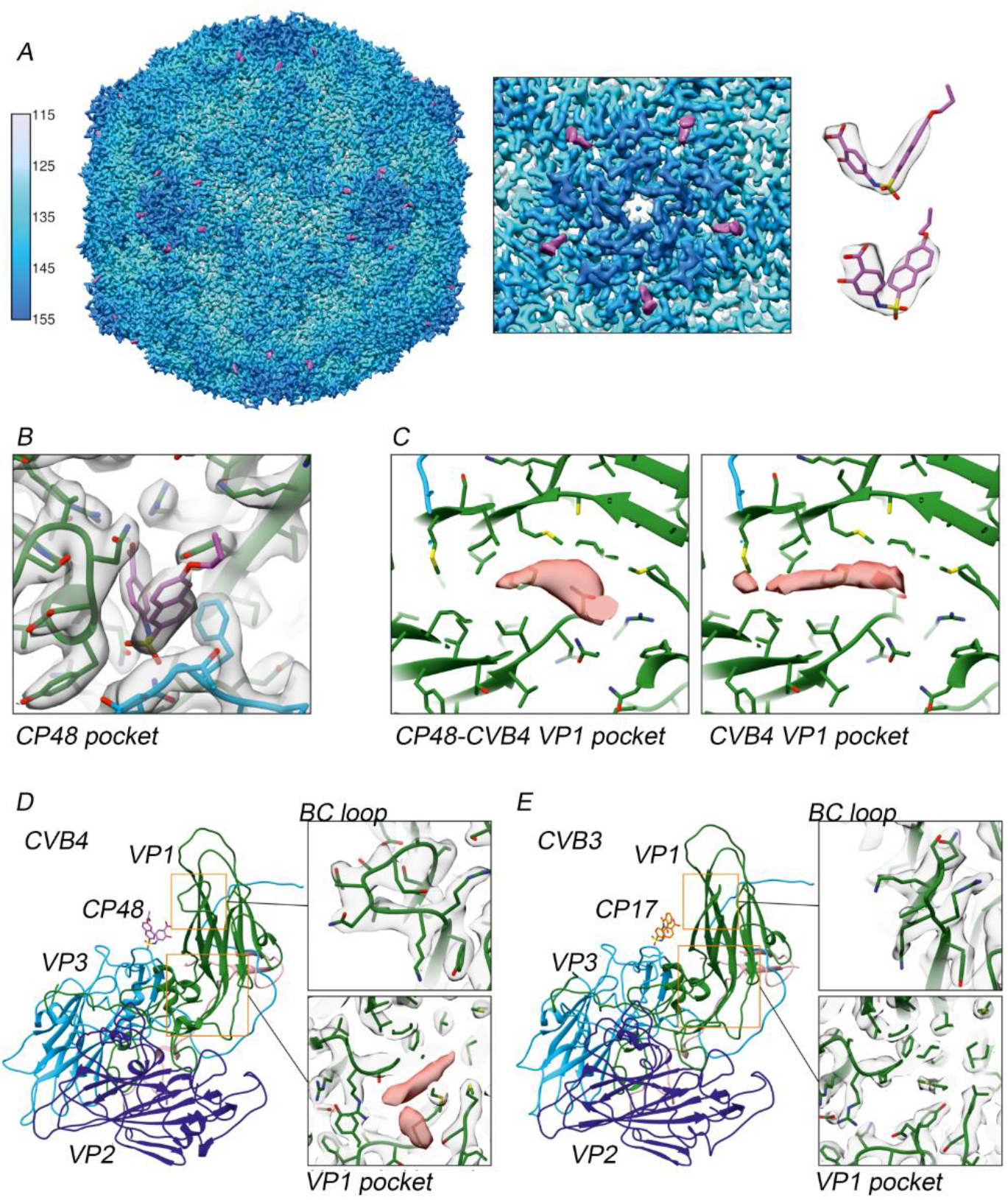
Cryo-EM of the CP48-CVB4 complex. (A) Visualization of CVB4 in the presence of CP48. The view is along the 2-fold axis with radial coloring, and the inset shows clear drug density at interprotomer site near the 5-fold axis of symmetry. Density for the 385.44 dalton CP48 is displayed in pink. (B) A close-up view of the capsid binder within the interprotomer pocket. (C) Density at the hydrophobic pocket of VP1 in CP48-CVB4 versus a control structure of CVB4 alone. (D-E) Two notable differences between CVB4 and CVB3 occur in the BC loop and pocket of VP1.

Aside from the interprotomer site, there were other notable differences in the CP48-CVB4 structure. First, in CP48-CVB4, density corresponding to the native pocket factor within the β-barrel of VP1 is altered by the presence of drug; however, there were no conformational changes in the four capsid proteins based on comparison to a control structure of CVB4 without CP48 (Figure 3C). Also, the control CVB4 structure has no density filling the interprotomer pocket. The alteration may result from a change in the structural dynamics of the pocket factor, or because this region is an additional weak binding site for CP48. The CVB3 Nancy strain lacks density for the lipid factor (Figure 3E). Furthermore, the Nancy strain has a substitution of Leu for Ile at position 92 in the hydrophobic pocket of VP1, which correlates with resistance to pleconaril-like compounds. Experiments involving a pleconaril-sensitive CVB3 Nancy variant (VP1_L92I) showed that when pleconaril and CP17 are combined, they have a synergistic effect, which suggests that the druggable site in VP1 does not contribute to the antiviral mechanism of interprotomer-targeting compounds (Abdelnabi et al., 2019). The second difference between CVB3 and CVB4 occurs in the exposed loop linking β-strands B and C of VP1, which for CVB4 differs in length and conformation (Figure 3D). This loop is known to be a serotype-specific neutralizing antigenic site (Reimann et al., 1991).

### Three key residues form a virion-stabilizing contact network in the presence of binders

Precise mechanistic descriptions of how capsid binders target enteroviruses has many practical applications, in terms of both understanding the cell biology of virus entry and design of new therapeutic agents. However, the large, flat, and relatively featureless surface of enterovirus capsids poses many challenges for drug design. Here, we used high-resolution cryo-EM to unveil a virion-stabilizing network that forms within the recently discovered druggable interprotomer pocket when an inhibitor is present. Although about 25 residues form the structural core of the binding site, our results indicate that a contact network of only three strictly conserved residues, each from a different polypeptide chain, dictate the size and shape of interprotomer-targeting compounds, as well as the mechanism of action. When an inhibitor is stably anchored to the network (60 sites per a single virion), it interferes with motion transmission such that the virus particle cannot undergo expansive conformational changes in the interprotomer region, and hence is unable to uncoat the genome at precisely the right time in infection. Interestingly, Duyvesteyn et al. recently published a 1.8 Å resolution X-ray structure of bovine enterovirus F3 (EV-F3) with glutathione (GSH) positioned in a similar way within the pocket (Figure 4A) (Duyvesteyn et al., 2020). The antioxidant engages the same virion-stabilizing network as CP17 and CP48, which is not surprising given that these molecules have strikingly similar geometrical and chemical features (Figure 4B and 4C). Specifically, GSH adopts a hook-shaped structure with a carboxylic end and a sulfur-containing elbow region. The overall size approximates that of inhibitors that occupy the interprotomer site. The carboxylic group interacts with the two Arg residues from neighboring VP1 polypeptides inside the pocket, while the sulfur atom interacts with the oxygen in the Gln side chain of VP3. It is believed that for CVB3 and CVB4, GSH makes strong interactions with adjacent protomers to facilitate intracellular assembly of progeny virions (Smith and Dawson, 2006, Ma et al., 2014, Thibaut et al., 2014). These residues that form the contact network motif within the interprotomer pocket are found across existing enterovirus structures (Figure 4D).

**Figure 4:**
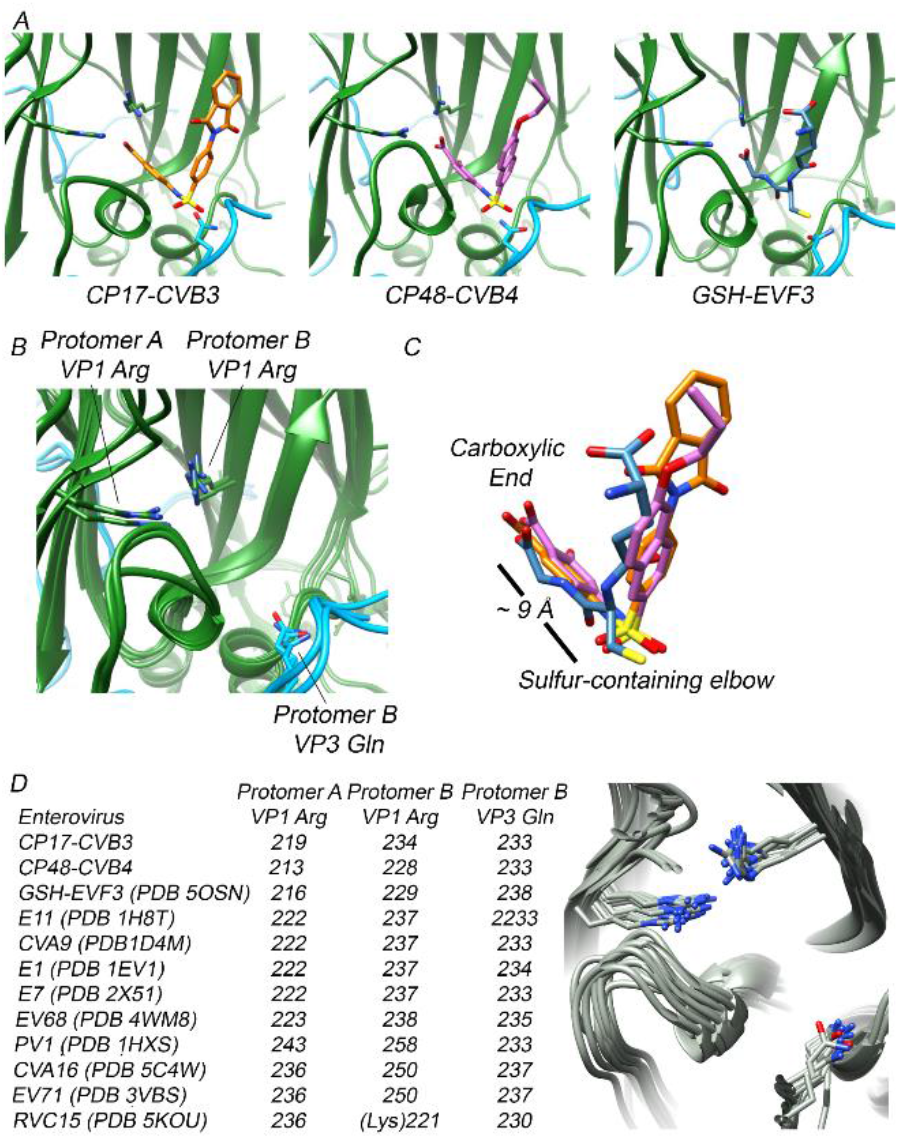
Stabilization inside the interprotomer pocket of enteroviruses. (A) Experimental structures of unique small molecules bound at interprotomer sites in CVB3, CVB4, and EV-F3 (PDB 5OSN). (B) Overlay of the three binding pocket residues, each from a different polypeptide chain, that confer capsid stabilization. (C) Interprotomer-targeting molecules have similar features. (D) Structural alignment of the three key residues that form the virion-stabilizing network inside the interprotomer pocket. The numbering of the residues in columns 2,3, and 4 were taken from the wwPDB files listed in column 1.

## Discussion

Despite decades of research on WIN antiviral compounds, no drugs have been approved for use against enteroviruses. Recently, a new class of broad-spectrum capsid binders was described, which inhibit a variety of enteroviruses by occupying a positively charged surface depression in the interprotomer zone. Structure-guided in vitro assays showed that pocket-bound molecules increase particle stability, however, issues related to resolution and mutational capacity (several of the CVB3 variants used in the study including VP1_R234G and VP3_Q233G were nonviable) limited characterization of drug-target interactions. In an effort to overcome the limitation, we have determined high-resolution cryo-EM structures of two enteroviruses complexed with interprotomer-targeting compounds. The quality of the maps was such that we could see clear densities for protein and ligand components, and modeling allowed us to visualize a conserved virion-stabilizing network within the pocket. The network extends from two Arg residues at the deepest part of the pocket to a Gln residue at the entrance via three connectors. Specifically, two Arg side chains from neighboring VP1 polypeptides interface with the carboxylic end of the inhibitor, and a third interaction occurs between the C-terminus of VP3 and the sulfur-containing elbow region of the inhibitor. The resulting effect of inhibitors secured to the network is a stabilization of several thousand kcal/mol for the virion. Aside from synthetic compounds, GSH targets and utilizes this stabilizing network to promote initial assembly of enteroviral protomers into pentamers. This finding, along with the structures reported here, indicate that three interprotomer residues, though not all in direct contact with each other, play a crucial role in forming stable virus capsids, and contextualize why this pocket is such an attractive drug target.

## Data and Code Availability

Coordinates and EM maps are available at the PDB and the Electron Microscopy Databank (EMDB): CVB4-CP48 PDB 6ZCK and EMD-11165, CVB3-CP17 PDB 6ZCL and EMD-11166, and CVB4 PDB 6ZMS and EMDB-11300.

## Acknowledgements

We thank Benita Löflund and Pasi Laurinmäki of Instruct-ERIC Center Finland and the Biocenter Finland National Cryo-EM microscopy unit, Helsinki University; Marta Carroni, the SciLifeLab Stockholm, and the CSC-IT Center for Science Ltd. for providing technical assistance and facilities to carry out the work. Additionally, we thank Johan Neyts and the Rega Institute for Medical Research in Leuven for kindly providing the CVB4 strain E2 used in this study. This project was supported by the Academy of Finland (315950 to S.J.B.), the Sigrid Juselius Foundation (to S.J.B.), and the People Programme (Marie Curie Actions) of the European Union’s Seventh Framework Programme FP7/2007-2013/ under REA grant agreement n°612308.

## Competing interest

None declared.

## Methods

### Virus culture and purification

BGM cells were cultured in Eagle’s minimum essential medium (MEM) supplemented with 10% fetal bovine serum (FBS), 1x nonessential amino acids, 1% GlutaMAX, and 1% antibiotic-antimycotic solution in a chamber environment adjusted to 37°C and 5% CO_2_. To produce virus particles for the study, 30 confluent T175 flask were inoculated with CVB4 (GenBank: AF311939.1) at a multiplicity of infection of ~0.5 in serum-free medium. Additionally, the infection medium contained 20 mM HEPES (pH 7.0). At 3 days post-infection, widespread viral cytopathic effect was evident, and the contents of each flask were collected, freeze-thawed 3 times, and centrifuged at 4,000 rpm and 4°C for 10 minutes to remove cellular debris. The supernatant was then carefully removed and concentrated using a Centricon centrifugal filter device (100 kilodalton cut-off). Virus particles were purified by centrifuging through a CsCl gradient (top density 1.25 g/cm^3^ and bottom density 1.48 g/cm^3^) at 30,000 rpm and 4°C for 19 hours. The gradient/exchange buffer consisted of 10 mM HEPES (pH 7.0), 150 mM NaCl, 2 mM MgCl_2_, and 2 mM CaCl_2_. Bands containing intact virions were collected and the CsCl was removed by buffer exchange.

### Thermostability assay

Approximately 5 × 10^4^ TCID_50_ units of CVB4 strain E2 was mixed with 20 μM concentration of CP48 in six tubes (reaction volume 52 μL) and incubated over a range from 37–52°C for 2 minutes, followed by rapid cooling on ice. The infectious virus load in the samples was estimated by an end-point titration assay using BGM cells. Specifically, serial 10 log dilutions were prepared in infection medium and applied to BGM cell monolayers on 96-well plates arranged one day before use by seeding 2 × 10^4^ cells per well. Two days after infection, the BGM cell monolayers were examined for cytopathic effects and TCID_50_/mL was estimated using Kärber-Spearman formula (Ramakrishnan, 2016).

### Cryo-EM sample preparation and data collection

The compound 4-{[(6-propoxy-2-naphthyl)sulfonyl]amino}benzoic acid (CP48) was ordered from a commercial supplier (www.specs.net) and dissolved in DMSO at a concentration of 10 mg/mL. Purified CVB4 and CP48 were then mixed at a molar ratio of 1:2,500 and incubated at 37°C for 1 hour. For cryo-EM sample preparation, 3.0 μL samples of CVB4-CP48 were applied to glow-discharged grids (Ted Pella product No. 01824). Grids were manually blotted with filter paper to remove excess sample and flash-frozen in liquid ethane with a homemade plunger. Data acquisition was carried out at the Swedish National Facility. The frozen-hydrated grids were loaded into and FEI Titan Krios electron microscope operated at 300 kV. A total of 4,379 movies were acquired with a Gatan K2 Summit direct electron detection camera at a nominal magnification of ×130,000, giving a pixel size of 1.06 Å per pixel. The total electron dose was approximately 46 electrons per Å^2^ fractionated into 30 frames. Frame images in each movie were aligned and averaged to correct for beam-induced motion using MotionCor2 (Li et al., 2013). A control dataset of CVB4 without CP48 was collected within the Electron Microscopy Core Unit at University of Helsinki using a Talos Arctica equipped with a Falcon III direct electron detection camera. Images were acquired at a nominal magnification of ×120,000, giving a pixel size of 1.24 Å per pixel. The accumulated electron dose was approximately 30 electrons per Å^2^ fractionated into 30 frames. MotionCor2 was used to produce a single micrograph from aligned and averaged movie frames.

### Image Processing

Defocus values of CVB4-CP48 micrographs were determined by Gctf (Zhang, 2016). A total of 31,436 particles were picked from 4,377 micrographs using ETHAN (Kivioja et al., 2000). Orientation and center parameters were determined and refined using RELION-3 within the Scipion image processing framework (Zivanov et al., 2018, de la Rosa-Trevin et al., 2016). Reference-free two-dimensional classification was used to discard 12,310 particles in poorly defined classes or false positives. An ab initio model generated with the RELION 3D Initial Model protocol was used as an initial reference model for maximum-likelihood three-dimensional classification. One class containing 18,626 high-quality particles was selected and divided into random halves for further refinement. The initial round of refinement was followed by subsequent rounds that included iterative per-particle CTF-refinement, and Bayesian polishing. After convergence in the final refinement step, and FSC curve was calculated and the resolution was determined to be 2.7 Å according to the gold-standard FSC = 0.143 threshold criterion. A B-factor of −70 Å^2^ was applied to sharpen the density map for modeling and analysis. The same image processing approach was applied to CVB4 without drug, as well as the raw data for CVB3-CP17 (EMPIAR-10199), which resulted in 3.4 Å and 2.8 Å maps, respectively. ResMap images, map cross-sections, and FSC curves for the three structures are included in Supplemental Figure 1 (Kucukelbir et al., 2014). A table for data collection, processing, and modeling (MolProbity) is included as Supplemental Table 1 (Chen et al., 2010).

### Modeling

An initial template for CVB4 capsid proteins VP1-VP4 was derived from a homology-based model calculated by I-TASSER (Yang et al., 2015). The UCSF Chimera Build Structure tool was used to translate the Simplified Molecular Input Line Entry Specification (SMILES) string for CP48 into a three-dimensional structure and parameterization was completed using SwissParam (Pettersen et al., 2004, Zoete et al., 2011). Structures for viral proteins and drug were docked into the EM density using UCSF Chimera, followed by iterative manual adjustment and real-space refinement using COOT (Emsley and Cowtan, 2004). Sequence assignment was guided by bulky amino acid residues such as Phe, Tyr, Trp and Arg, and featureful density allowed placement of the ligand. The optimized model for CVB4-CP48 was then subjected to end-stage refinement using the Molecular Dynamics Flexible Fitting (MDFF) program originally developed by Klaus Schulten and coworkers (Trabuco et al., 2008). Harmonic restraints were applied to prevent overfitting during simulations. Capsid proteins for the CVB4 virion without drug were modeled using a similar protocol and comparison to the CP48-CVB4 structure confirmed that drug binding does not induce conformational changes in the virion. The RMSD between CP48-CVB4 and CVB4 alone was 0.45. The binding energy for CP48 inside the interprotomer pocket was obtained using the NAMD energy plugin in VMD (Phillips et al., 2005, Humphrey et al., 1996). We used the same procedure to refine atomic coordinates for CVB3-CP17 (PDB ID code 6GZV) into the newly determined CVB3-CP17 2.8 Å map. The structural alignment of PDB files was done using the MatchMaker feature of UCSF Chimera.

---

**Supplementary Figure 1:**
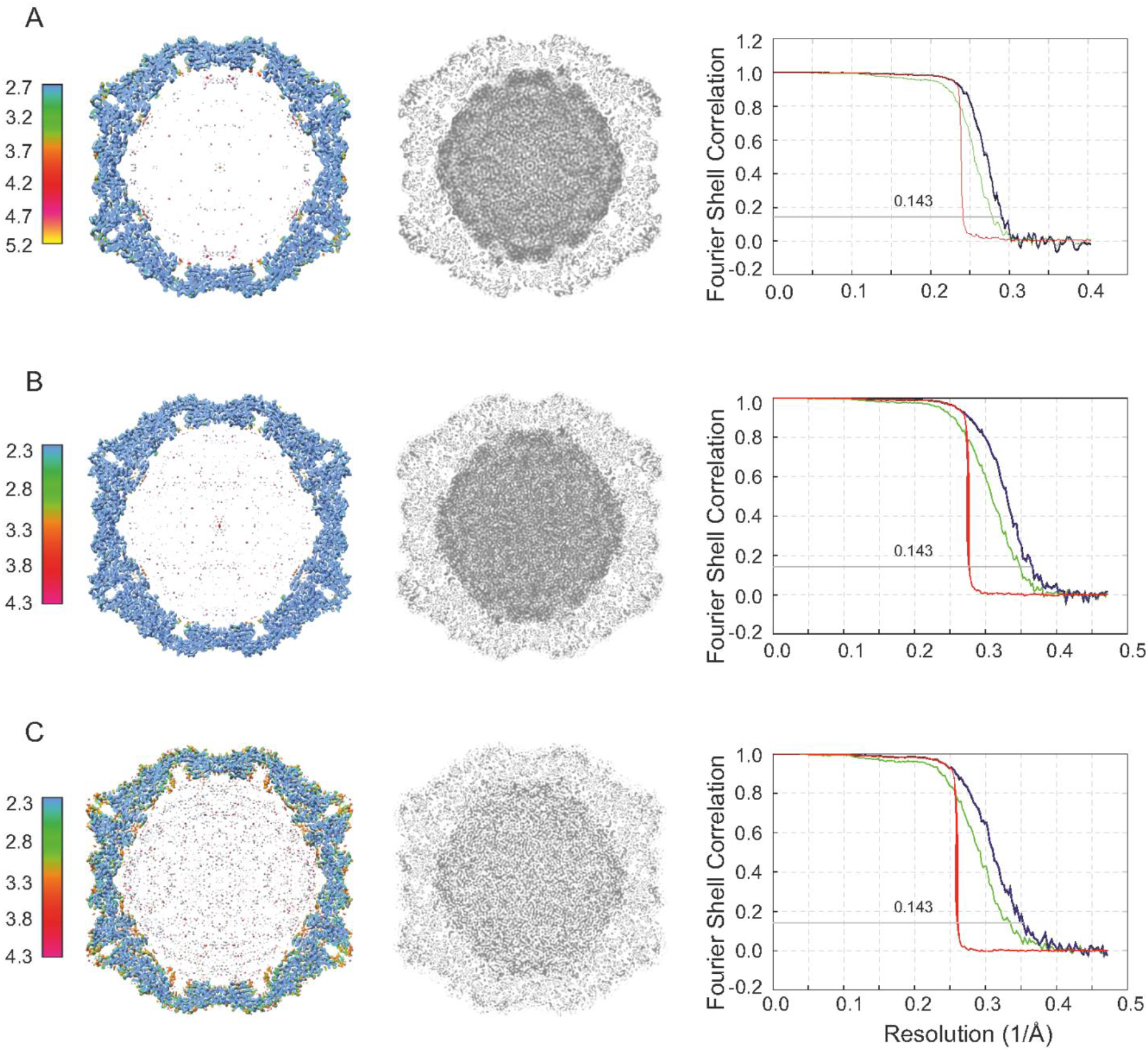
ResMap resolution slices (left), central cross-sections (middle), and FSC plots (right) are shown for CVB4, CVB4-CP48, and CVB3-CP17.

**Supplementary Table 1:** Cryo-EM data collection, processing, and modeling.

| **Data collection and processing** | **CP48-CVB4** | **CP17-CVB3** | **CVB4** |
| --- | --- | --- | --- |
| Magnification | 130,000 | 130,000 | 120,000 |
| Voltage | 300 | 300 | 200 |
| Electron exposure (e-/Å<sup>2</sup>) | 47 | 47 | 30 |
| Defocus range (µM) | -1 to -5 | -1 to -5 | -1 to -5 |
| Pixel Size (Å) | 1.06 | 1.06 | 1.24 |
| Symmetry imposed | I2 | I2 | I2 |
| Final particle images | 13,252 | 18,626 | 40,627 |
| Map resolution (Å) | 2.7 | 2.8 | 3.4 |
| FSC threshold | 0.143 | 0.143 | 0.143 |
| **Refinement** |  |  |  |
| Map sharpening B factor (Å<sup>2</sup>) | -70 | -77 | -90 |
| Model resolution | 2.7 | 2.8 | 3.4 |
| Protein residues | 800 | 798 | 800 |
| Ligands | 1 | 1 | 0 |
| Myristoylation | 1 | 1 | not modeled |
| Lipid factor (VP1) | not modeled | 0 | not modeled |
| MolProbity score | 1.40 (97 percentile) | 1.16 (99th percentile) | 1.52 (95 percentile) |
| Clashscore | 0 | 0 | 0 |
| Poor rotamers | 25 (3.9%) | 15 (2.4%) | 11 (1.7%) |
| Favored rotamers | 588 (91%) | 660 (94%) | 615 (95%) |
| Ramachandran outliers | 13 (1.7%) | 9 (1.1%) | 23 (3.1%) |
| Ramachandran favored | 701 (93%) | 774 (94%) | 686 (91%) |

## Notes

### Competing Interest Statement

The authors have declared no competing interest.

